# A *Vibrio cholerae* BolA-like protein is required for proper cell shape and cell envelope integrity

**DOI:** 10.1101/597369

**Authors:** Aurore Fleurie, Abdelrahim Zoued, Laura Alvarez, Kelly M. Hines, Felipe Cava, Libin Xu, Brigid M. Davis, Matthew K. Waldor

**Affiliations:** Department of Microbiology, Harvard Medical School, Boston, MA 02115, USA.; Division of Infectious Diseases, Brigham and Women’s Hospital, Boston, MA 02115, USA.; Laboratory for Molecular Infection Medicine Sweden, Department of Molecular Biology, Umeå Centre for Microbial Research, Umeå University, Umeå SE-90187, Sweden; Department of Medicinal Chemistry, University of Washington, Seattle, WA 98195, USA.; Howard Hughes Medical Institute, Boston, MA 02115, USA.

**Keywords:** IbaG, BolA, *Vibrio cholerae*, cell shape, cell envelope, iron-sulfur cluster

## Abstract

BolA family proteins are conserved in gram-negative bacteria and many eukaryotes. While diverse cellular phenotypes have been linked to this protein family, the molecular pathways through which these proteins mediate their effects are not well-described. Here, we investigated the role of BolA family proteins in *Vibrio cholerae*, the cholera pathogen. Like *Escherichia coli*, *V. cholerae* encodes two BolA proteins, BolA and IbaG. However, in marked contrast to *E. coli*, where *bolA* is linked to cell shape and *ibaG* is not, in *V. cholerae, bolA* mutants lack morphological defects, whereas *ibaG* proved critical for the generation and/or maintenance of the pathogen’s morphology. Notably, the bizarre-shaped, multi-polar, elongated and wide cells that predominated in exponential phase Δ*ibaG V. cholerae* cultures were not observed in stationary phase cultures. The *V. cholerae* Δ*ibaG* mutant exhibited increased sensitivity to cell envelope stressors, including cell wall acting antibiotics and bile, and was defective in intestinal colonization. Δ*ibaG V. cholerae* had reduced peptidoglycan and lipid II and altered outer membrane lipids, likely contributing to the mutant’s morphological defects and sensitivity to envelope stressors. Transposon-insertion sequencing analysis of *ibaG’*s genetic interactions suggested that *ibaG* is involved in several processes involved in the generation and homeostasis of the cell envelope. Furthermore, co-purification studies revealed that IbaG interacts with proteins containing iron-sulfur clusters or involved in their assembly. Collectively, our findings suggest that *V. cholerae* IbaG controls cell morphology and cell envelope integrity through its role in biogenesis or trafficking of iron-sulfur cluster proteins.

**Importance:** BolA-like proteins are conserved across prokaryotes and eukaryotes. These proteins have been linked to a variety of phenotypes, but the pathways and mechanisms through which they act have not been extensively characterized. Here, we unraveled the role of the BolA-like protein IbaG in the cholera pathogen *Vibrio cholerae*. The absence of IbaG was associated with dramatic changes in cell morphology, sensitivity to envelope stressors, and intestinal colonization defects. IbaG was found to be required for biogenesis of several components of the *V. cholerae* cell envelope and to interact with numerous iron-sulfur cluster containing proteins and factors involved in their assembly. Thus, our findings suggest that IbaG governs *V. cholerae* cell shape and cell envelope homeostasis through its effects on iron-sulfur proteins and associated pathways. The diversity of processes involving iron-sulfur containing proteins is likely a factor underlying the range of phenotypes associated with BolA family proteins.

## Introduction

The BolA protein family is widely conserved across gram-negative bacteria and eukaryotes (1). These proteins have been linked to a range of cellular phenotypes, including cell morphology, membrane permeability, motility, and biofilm formation (2). BolA-like proteins have a class II KH fold related to that of the OsmC hyperperoxide reductase (3) that includes a helix-turn helix domain (HTH) (1). The HTH domains of certain BolA proteins have been shown to bind DNA and modulate transcription (4). In several species, BolA family members have been linked to stress response pathways, and the absence or overexpression of BolA proteins can modulate bacterial viability in response to a variety of environmental challenges. Also, there is an emerging understanding of a role for BolA proteins in iron homeostasis and iron-sulfur cluster assembly and trafficking (5). The varied genomic context for genes encoding BolA family members, the range of phenotypes associated with BolA proteins, and the fact that some organisms encode multiple BolA family members suggests that these proteins likely contribute to a variety of processes, both across species and within a single species. However, mechanisms underlying these proteins’ diverse effects on cell physiology have largely not been determined.

In *E. coli*, *bolA* expression is induced in response to several stressors (6). Overexpression of *bolA* in *E. coli* induces formation of spherical cells (7), potentially due to associated upregulation of *dacA* and *dacC*, which encode the penicillin binding proteins (PBPs) PBP5 and PBP6, as well as to downregulation of *mreB* (8–10). Overexpression of *bolA* is also thought to decrease the permeability of the bacterial outer membrane, while the absence of *bolA* alters the accessibility of outer membrane proteins (11). Transcriptomic and ChIP analyses have shown that BolA overexpression directly modulates transcription (4). Finally, a role for BolA in biofilm formation has been demonstrated (4, 12, 13).

*E. coli*, like many other organisms, encodes more than one BolA family protein. In addition to the 105 amino acid protein BolA*, E. coli* encodes IbaG (formerly YrbA), an 84 amino acid protein that also contains the characteristic class II KH fold of BolA proteins (14). Unlike BolA, neither overexpression nor the absence of IbaG alters *E. coli* cell shape; however, overexpression of *ibaG* is deleterious to bacterial growth, while its deletion enhanced bacterial growth (14). *IbaG* expression is induced in response to acid, accounting for its name, **I**nfluenced **b**y **a**cid **g**ene, and the absence of *ibaG* also increases *E. coli* sensitivity to acid stress. Although IbaG, like BolA, is presumed to act as a transcription factor, it does not appear to recognize sequences bound by BolA (14). Thus, in *E. coli*, IbaG’s role is distinct from that of BolA.

BolA-like proteins have roles in genesis of iron-sulfur proteins through their partnerships with monothiol glutaredoxins (Grxs). Bioinformatics analysis of co-occurrence provided the first clue linking Grx proteins and BolA-like proteins; the simultaneous presence or absence in many genomes of genes encoding both these proteins suggested a functional interaction between them (1). Subsequently, it has been shown in *E. coli* and several eukaryotes that monothiol Grxs and BolA proteins form heterocomplexes implicated in iron-sulfur cluster assembly and trafficking (5). In particular, *E. coli*’s single monothiol Grx (Grx4) forms [2FE-2S]-bridged heterodimers with BolA and IbaG (15, 16). Both *grxD* (encoding Grx4) and *ibaG* have also been found to exhibit aggravating genetic interactions with genes in the *isc* operon, which encodes components of the housekeeping iron-sulfur cluster assembly pathway, suggesting that Grx4 and IbaG may mediate an alternate process of iron-sulfur cluster assembly (17).

Like *E. coli*, the gram-negative pathogen *V. cholerae* encodes two members of the BolA protein family, BolA and IbaG. *IbaG* has a similar genomic context in both organisms; it is found downstream of *mlaBCDEF*, which encode components of an ABC transport system required for maintenance of outer membrane lipid asymmetry, and upstream of *murA*, whose product catalyzes the first step in peptidoglycan assembly (Fig. 1A). In contrast, genomic placement of *bolA* is not conserved between *V. cholerae* and *E. coli*. To date, no role has been reported for either BolA family member in *V. cholerae*.

**Figure 1.**
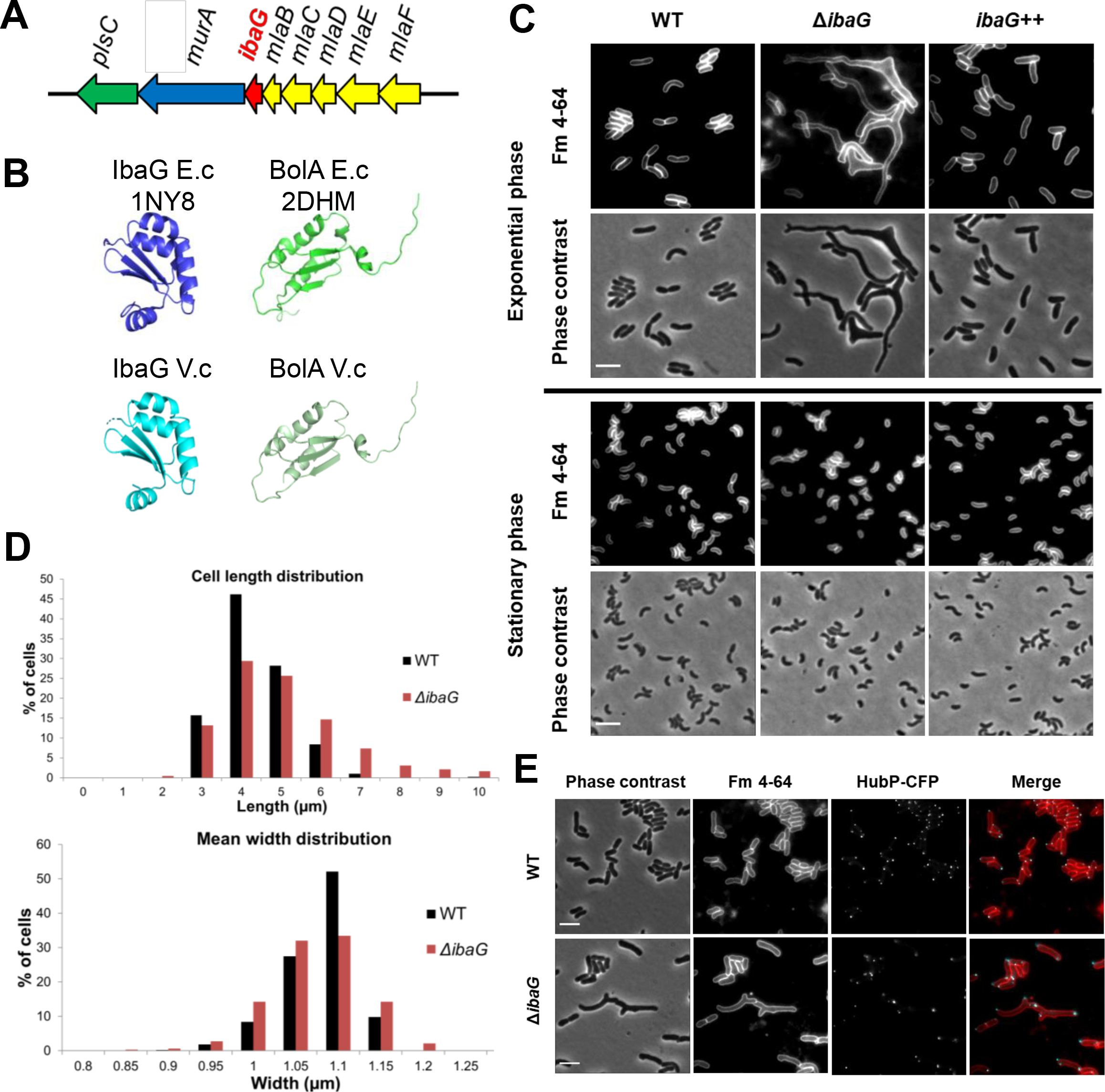
IbaG is required for *Vibrio cholerae* cell shape. **A**. Schematic of the genomic neighborhood of *ibaG* (red), which includes *mla* BCDEF (yellow), implicated in maintenance of outer membrane lipid asymmetry, *murA* (blue), involved in synthesis of peptidoglycan precursors, and *plsC* (green), implicated in phospholipid synthesis. **B**. Comparison of structures of BolA and IbaG in *E. coli* and *V. cholerae*. Structure of IbaG (PDB accession code 1NY8, left) and BolA from *E. coli* (E.c) (PDB accession code 2DHM, right) (upper panel) and the predicted models obtained with PHYRE2 for IbaG and BolA from *V. cholerae* (V.c) (lower panel). **C**. Phase contrast and fluorescence imaging of FM4-64-stained WT, Δ*ibaG*, and *ibaG* overexpressing (*ibaG*++) *V. cholerae* grown to exponential and stationary phase in M9 medium. Scale bars 2 μm. **D**. Cell length and mean width distribution of WT and Δ*ibaG* strains grown in M9 medium. At least 1000 cells were measured for each condition using MicrobeTracker. Statistical significance was determined using a non-parametric Mann Whitney U test. P-value ≤0.001. **E**. Fluorescence imaging of WT and Δ*ibaG* strains grown in M9 medium and expressing a chromosome-encoded HubP-CFP. Cells were also stained with FM4-64. Scale bars 2 μm.

Here, we explored the role of BolA family proteins in *V. cholerae*, the cholera pathogen. In marked contrast to *E. coli*, we found that *V. cholerae ibaG* is critical for the generation and/or maintenance of the pathogen’s morphology. Δ*ibaG V. cholerae* exhibited increased sensitivity to cell envelope stressors and were defective in intestinal colonization. These defects are likely attributable to the aberrant composition of the mutant’s cell envelope, including reduced peptidoglycan and altered outer membrane lipids. Genetic and protein interaction analyses suggest that IbaG may control *V. cholerae* cell morphology and envelope integrity through its role in biogenesis or trafficking of iron-sulfur cluster proteins.

## Results

### IbaG is required for *V. cholerae* cell morphology and growth

The predicted amino acid sequences of BolA and IbaG in *V. cholerae* and *E. coli* are highly similar (Fig. S1AB). Furthermore, the predicted structures of homologous proteins are nearly identical for the two species (Fig. 1B). In contrast, the two *V. cholerae* BolA-family proteins only share 24% amino acid similarity despite the conservation of their secondary structures (Fig. S1CD). We constructed derivatives of *V. cholerae* N16961 in which either *ibaG* or *bolA* is deleted or overexpressed. Phase contrast and fluorescence microscopy of these cells revealed no effect of *bolA* on cell shape (Fig. S2A). In marked contrast, exponential phase Δ*ibaG* cells had grossly distorted cell shapes (Fig. 1CD), whereas overexpression of *ibaG* did not influence *V. cholerae* morphology (Fig. 1CD). The morphological defects of the *ibaG* deletion mutant were observed in both LB and M9 media but were more pronounced in the latter (Fig. S2B). The Δ*ibaG* cells were generally longer and wider than the wild-type (WT); moreover, many of the mutant cells exhibited branching and the presence of extra cell poles (Fig. 1CDE). HubP, a key regulator of *V. cholerae* pole development (18), was often detectable at all poles (Fig. 1E), suggesting that the polar cell domain is intact at the supernumerary poles in branched Δ*ibaG* cells.

Notably, the mutant’s morphological defects were only observed during exponential phase growth; in stationary phase, Δ*ibaG* cells exhibited normal shape and size (Fig. 1C and Fig. S2C). These differences cannot be explained by changes in *ibaG* expression, which were very similar during exponential and stationary phase (Fig. S3A). The Δ*ibaG* morphological defects were eliminated by expression of *ibaG* in trans, indicating that shape changes are specifically linked to the absence of IbaG and not due to polar effects on other genes in the putative *ibaG* operon (Fig. S3BC). Thus, in marked contrast to *E. coli*, *ibaG* has a pronounced influence on *V. cholerae* morphology; furthermore, *bolA* does not appear to modulate *V. cholerae* cell shape, whereas its over-expression in *E. coli* results in shape defects (7, 8). Based on these observations, additional studies were focused on deciphering the role(s) of *ibaG* in *V. cholerae*.

Growth analyses of Δ*ibaG* and WT *V. cholerae* revealed that the deletion markedly reduced the growth rate and terminal density of cells cultured in M9 medium (Fig. 2A). In LB medium, the effect was much less dramatic; the terminal densities of WT and Δ*ibaG* cultures were equivalent, but the mutant strain had a prolonged lag phase. The impaired growth of Δ*ibaG V. cholerae* contrasts with that of Δ*ibaG E. coli*, which displays enhanced growth (14), providing additional evidence that *ibaG* plays distinct roles in these organisms.

**Figure 2.**
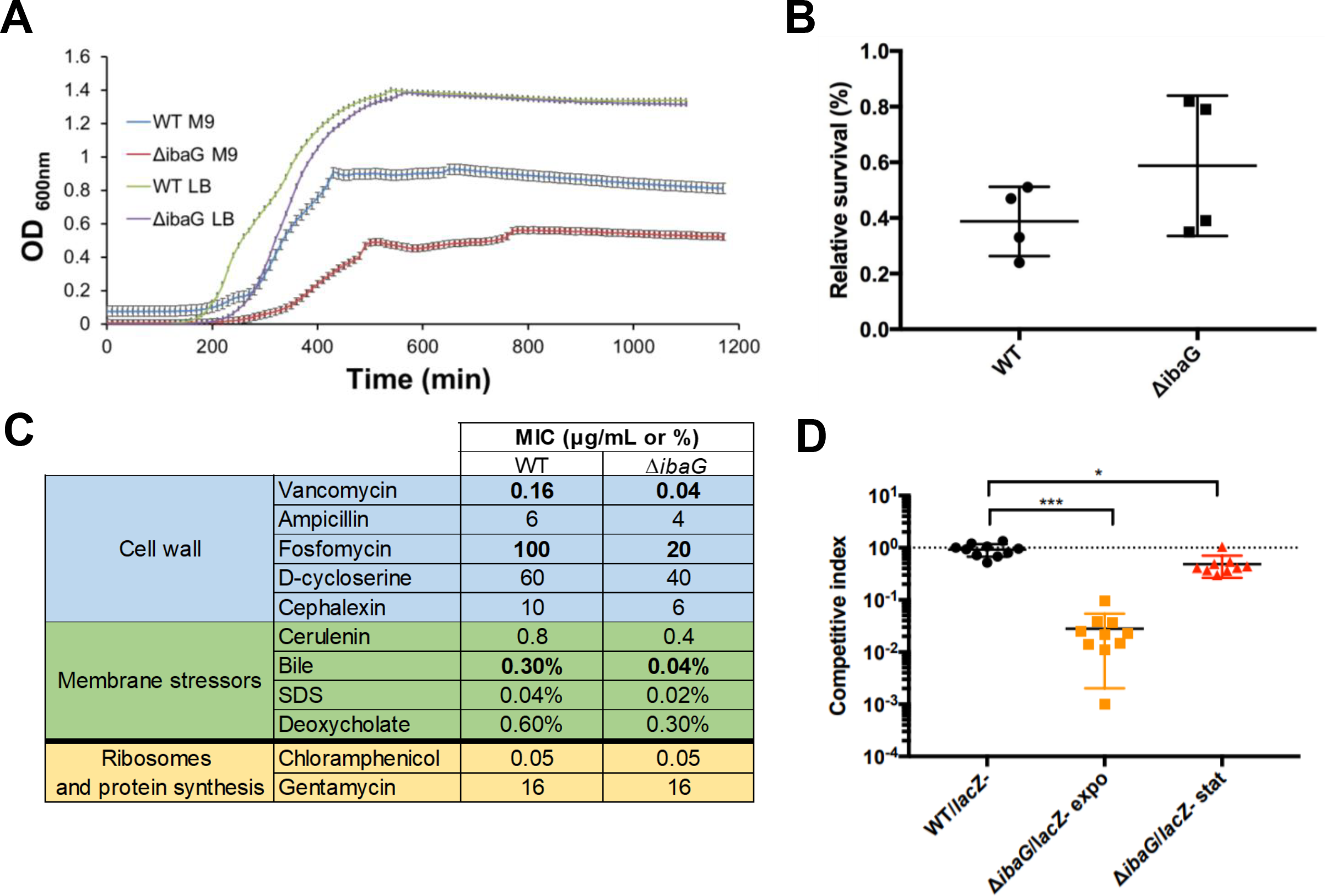
IbaG augments *V. cholerae* resistance to cell envelope stressors and promotes intestinal colonization. **A**. Growth curves of WT and Δ*ibaG V. cholerae* grown in M9 medium. OD_600nm_ was measured at 10-min intervals. Experiments were done in biological triplicate; error bars show standard deviations. **B**. Acid resistance was determined by calculating the proportion of cells that survived during growth in acidic medium (LB pH5.5) vs in LB pH7. Colony forming units/mL (CFU/mL) were determined for WT and Δ*ibaG* strains after one hour growth in acidic medium from exponential phase cultures. Experiments were performed in quadruplicate. **C**. Minimum inhibitory concentration (MIC) for indicated agents were measured after 24h growth in M9 at 37°C without shaking. The values shown represent the mean value obtained with two biological replicates done in technical quadruplicates for each strain. The concentrations are in µg/mL, except for bile, SDS and deoxycholate which are in %. **D**. Competitive indices for intestinal colonization for indicated strain pairs. Suckling mice were inoculated with 1:1 mixture of Δ*ibaG* and a *lacZ*-negative derivative of WT, made either from log phase (OD_600nm_ = ∼0.2) or overnight cultures. Competitive indices represent the output ratio (mutant strain CFU/*lacZ-* strain CFU) divided by the input ratio. Black horizontal lines represent geometric means and colored horizontal lines show standard deviation. A Mann Whitney U non-parametric test was used to assess statistical significance. *** =p-value ≤0.0001, * =p-value = 0.014

### IbaG promotes *V. cholerae* survival in the presence of factors that target the cell envelope

Since BolA family proteins have been shown to participate in stress response pathways (2), we explored whether the absence of *ibaG* altered *V. cholerae* survival following exposure to a range of environmental stresses. Given results from studies of *E. coli* IbaG, we first assessed whether *V. cholerae* IbaG modulates growth or survival under acidic conditions. Using a previously described acid resistance assay (19), we observed that the percentage survival of WT and ∆*ibaG* cells does not differ following a one-hour incubation in LB at pH5.5 (Fig. 2B). Additionally, we found that the growth rate of WT and ∆*ibaG V. cholerae* in LB pH5.5 are very similar, although a longer lag period prior to growth was evident for the ∆*ibaG* cells (Fig. S3D). Furthermore, quantitative RT-PCR analysis revealed no change in *ibaG* expression following a one hour exposure to acidified media (pH5.5) (Figure S3A). Thus, in contrast to *E. coli ibaG*, *V. cholerae ibaG* does not appear to promote bacterial resistance to acidic growth conditions.

To further evaluate the effect of IbaG on *V. cholerae* resistance to stressors, we determined the minimum inhibitory concentrations (MICs) of a wide variety of antimicrobial compounds for WT and *ΔibaG* cells. Notably, we observed that MICs for several antibiotics that target the cell wall (vancomycin, ampicillin, D-cycloserine, fosfomycin, cephalexin) were lower for the *ΔibaG* cells than for the WT (Fig. 2C). Additionally, we found that *ΔibaG* cells have increased sensitivity to bile, deoxycholate, SDS, and cerulenin (an inhibitor of fatty acid synthesis), all of which disrupt the outer membrane. In contrast, MICs of antibiotics that target the ribosomes and protein synthesis (chloramphenicol and gentamycin) were identical for the WT and the deletion strain. Collectively, these results suggest that the cell envelope of *ΔibaG V. cholerae* is more susceptible to disruption than that of WT cells, raising the possibility that IbaG regulates expression and/or the activity of factors that contribute to envelope production or maintenance.

### Exponential phase *ibaG V. cholerae* exhibit defective intestinal colonization

Given the sensitivity of the *ΔibaG V. cholerae* to cell envelope stressors including bile, we investigated whether *ibaG* contributes to the pathogen’s capacity to survive and proliferate in the intestines of suckling mice, a well-established model for studying *V. cholerae* intestinal colonization (20). Mice were orogastrically inoculated with 1:1 mixtures of WT and *ΔibaG* cells, and the relative abundance of the mutant cells within the small intestine was assessed at ~24 hours post-infection. Unexpectedly, the resulting competitive index for the *ΔibaG* cells was dependent upon the growth phase of cells used for the inoculum (Fig. 2D). When mice were infected with cells from stationary phase cultures, colonization by the *ΔibaG* cells was minimally attenuated. In contrast, when mice were infected with cells from log-phase cultures, the *ΔibaG* cells exhibited a ~50× decrease in colonization relative to the WT cells (Fig. 2D).

### *ΔibaG* cells have reduced peptidoglycan and phosphatidylethanolamine

Our observation that *ΔibaG* cells are more sensitive than WT *V. cholerae* to agents that disrupt the cell wall or the outer membrane prompted us to further explore the structure and composition of these components in the *ΔibaG* background. Peptidoglycan (PG) was isolated from exponential phase WT and *ΔibaG* cells and its abundance and composition were measured. *ΔibaG* cells contained ~25% less PG than did WT cells (Fig. 3A and S4). Furthermore, there were differences in the abundance of several PG constituents (Fig. 3B). In particular, PG from *ΔibaG* cells had shorter average chain lengths, and it contained more than twice the WT level of Lpp, an outer membrane protein that is covalently linked to PG and helps anchor it to the outer membrane. Although the precise consequences of these changes are difficult to predict, it is likely that the reductions and alterations in *ΔibaG* PG contribute to the increased sensitivity of these cells to antibiotics that interfere with PG synthesis.

**Figure 3.**
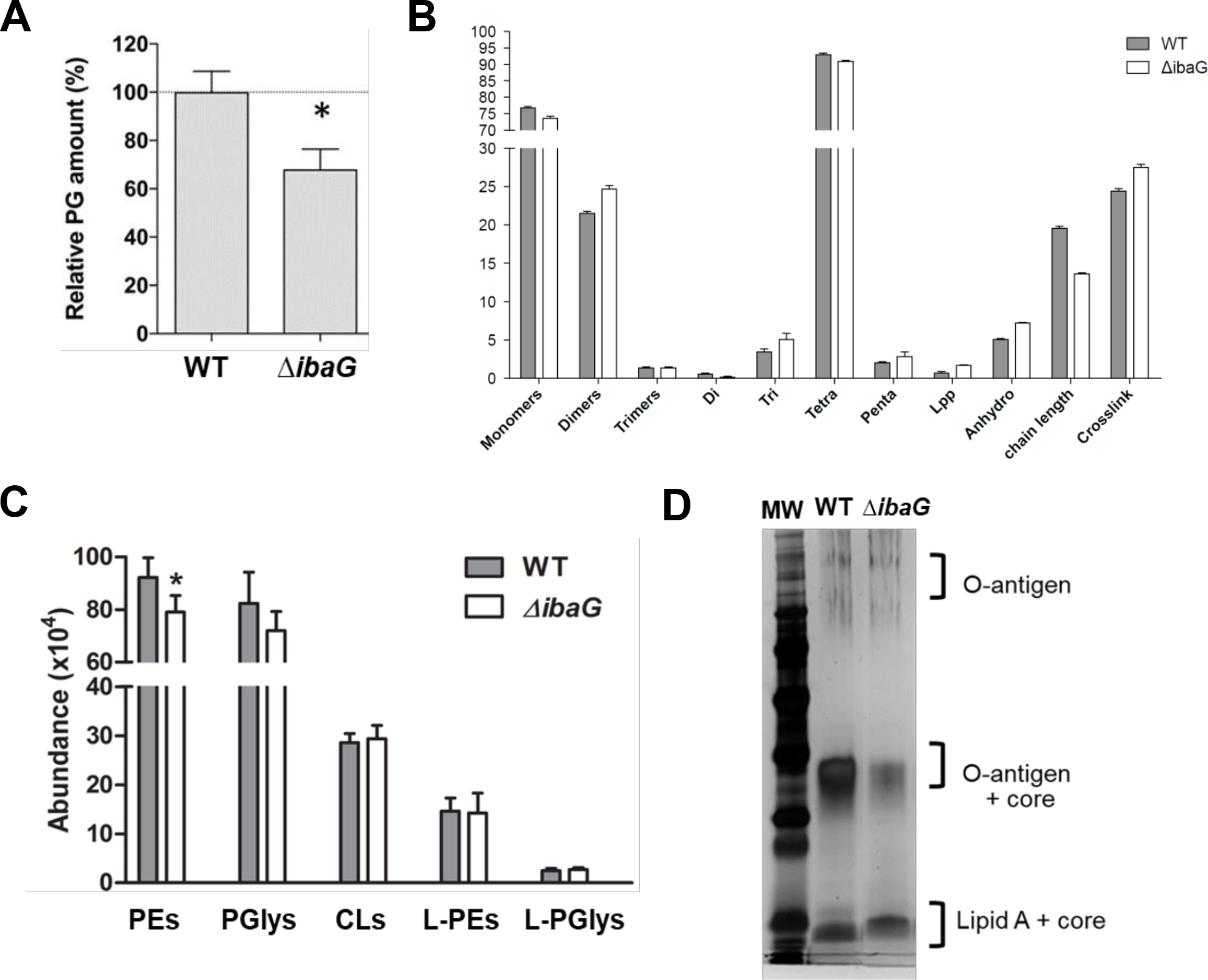
Comparison of cell envelope components in WT and *ΔibaG V. cholerae*. **A, B**. Abundance (A) and composition (B) of peptidoglycan (PG) isolated from exponential phase WT and *ΔibaG V. cholerae*. PG from each strain was analyzed in triplicate. Monomers, dimers and trimers represent muropeptides and di (dimers), tri (trimers), tetra (tetramers) and penta (pentamers) represent peptides; *= p-value <0.01 (t-test) **C**. Quantification of different lengths and saturated forms of phosphatidylethanolamines (PEs), phosphatidylglycerols (PGlys), cardiolipins (CLs), lyso-PEs (L-PE) and lyso-PGs (L-PGlys) in whole cell pellets from exponential phase WT and *ΔibaG V. cholerae*. *= p-value <0.01 (t-test) **D**. Silver stained SDS-PAGE of LPS isolated from exponential phase from WT and *ΔibaG* strains.

Lipidomic analysis, using hydrophilic interaction liquid chromatography-ion mobility-mass spectrometry (HILIC-IM-MS) of exponential phase WT and *ΔibaG* cells was also performed. The most obvious difference between the WT and Δ*ibaG* strains was an overall decrease in phosphatidylethanolamine (PE) (Fig. 3C), whereas phosphatidylglycerol (PGly) had a trend toward decrease and cardiolipin abundance was not significantly affected in the Δ*ibaG* strain (Fig. 3C). Since the biogenesis of 3-deoxy-d-manno-octulosonic acid (Kdo) requires PE (21, 22), and PE deficiency downregulates LPS biosynthesis in *E. coli* (21), we also quantified lipopolysacchararide (LPS) present in exponential phase WT and mutant cells. The Δ*ibaG* strain contained significantly less LPS than the WT strain (Fig. 3D). Decreased LPS levels might contribute to the Δ*ibaG* mutant’s increased sensitivity to membrane-disrupting factors such as bile and SDS and contribute to its colonization defect.

### Transposon insertion site-sequencing analysis links IbaG to cell envelope biogenesis

To gain further insight into the pathways and processes affected by *ibaG*, we carried out a comparative transposon-insertion sequencing analysis to identify transposon insertions that are underrepresented in the Δ*ibaG* background relative to the WT strain. Loci for which fewer insertions are identified in the Δ*ibaG* vs WT background are candidates for synthetic lethality with *ibaG* and may contribute to processes that are also impaired by the absence of *ibaG*. We identified 38 genes that were underrepresented at least two-fold in the Δ*ibaG* insertion library with a P value of < 0.05 (Fig. 4AB). Notably, more than 1/3 of these loci are involved in pathways linked to cell envelope integrity and/or LPS and PG synthesis (Fig. 4B). They include *mlaBCD*, which along with *mlaA* encode an ABC transport system involved in maintaining outer-membrane lipid asymmetry, PG biosynthetic gene *pbp1A* and its activator *lpoA*, several loci in the *rfa* cluster, which contains many of the genes for LPS synthesis, and genes encoding components of the *tol* system, which regulates PBP1B and is important for outer membrane stability (23) (Fig. 4C). Collectively, these results provide further support for the idea that *ibaG* is important for biogenesis and/or maintenance of the cell envelope, so that Δ*ibaG* cells are particularly sensitive to additional mutations that affect this structure.

**Figure 4.**
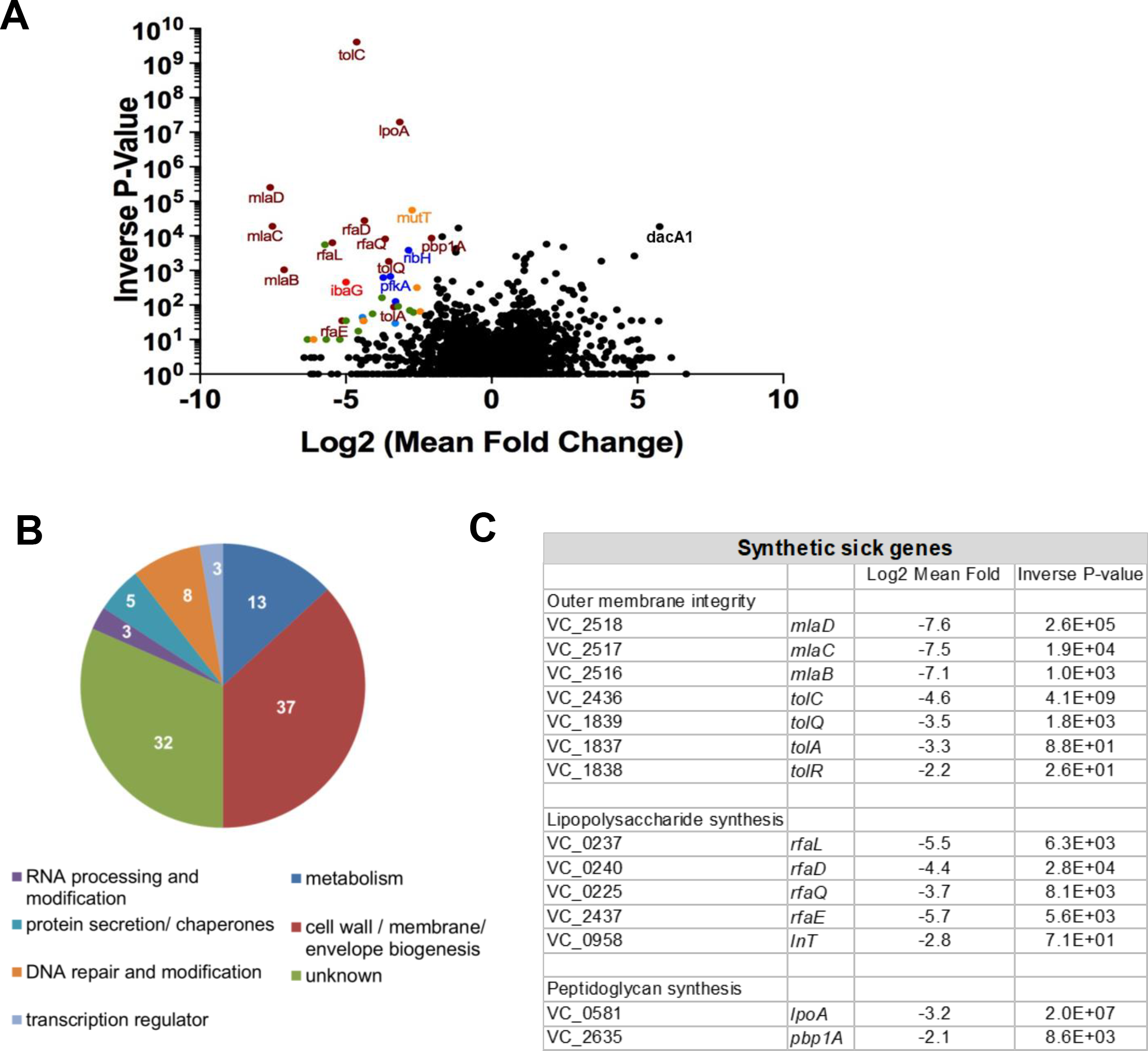
Transposon-insertion sequenced-based analyses of *ibaG* genetic interactions. **A**. Volcano plots depicting the relative abundance of read counts mapped to individual genes in transposon libraries made in Δ*ibaG* vs WT. For each gene, the Log2 mean fold change (*x* axis) and associated p*-*value (*y* axis) is shown. Genes shown in colors are considered significantly under-represented compared to the WT (mean fold change >2 and p-value <0.05) and the colors correspond to the functional classification represented in **B**. A comprehensive list of the genes over or under-represented in the Δ*ibaG* library with a mean fold change >2 and a p-value <0.05 is shown in Table S1. **B**. Functional classification of the genes classified as under-represented in the Δ*ibaG* background. Numbers represent the percentage of genes (of 38 total) in each category. **C**. Under-represented genes in the Δ*ibaG* insertion library that are involved in cell envelope integrity and/or LPS and peptidoglycan synthesis.

We also identified 34 loci that are overrepresented at least two-fold in the Δ*ibaG* insertion library with a P value of < 0.05 (Fig. 4A; Table S1). Intriguingly, these included *dacA1 (pbp5*), which encodes a low-molecular weight PG binding protein, which has been found to be necessary for normal *V. cholerae* growth and morphology (24). *V. cholerae* Δ*dacA*1 cells exhibited branches and aberrant poles and are wider as well as elongated (24), phenotypes which are strikingly reminiscent of the morphology of Δ*ibaG* cells. Disruption of *dacA1* also impedes *V. cholerae* cell growth and viability. However, in the *ibaG* background, the effects of *dacA1* disruption may be less detrimental, perhaps because they affect processes that have already been disrupted.

### IbaG interacts with numerous [Fe-S] cluster proteins

In addition to the genetic interactions revealed by transposon-insertion sequencing, we also identified proteins that interact with IbaG. In *E. coli*, IbaG interacts with Grx4, forming [2Fe-2S]-bridged heterodimers (16). Bacterial two-hybrid analysis demonstrated that the *V. cholerae* versions of these proteins also interact (Fig. S5). To further our knowledge of IbaG’s partners, epitope tagged IbaG was affinity purified, and co-purified proteins were identified *via* tandem mass spectrometry analysis (Fig. 5A, Table S2). Notably, a third of the proteins that copurified with IbaG have roles in either iron-sulfur cluster biogenesis (e.g. IscS, IscU), use iron-sulfur clusters as cofactors (e.g. NqrF, IspG), or bind iron-sulfur clusters and serve as carriers to transfer them to other proteins (e.g. NfuA, ErpA) (Fig. 5B). These interactions suggest that *V. cholerae* IbaG contributes to iron trafficking and can bind [Fe-S] clusters as shown for *E. coli* IbaG (16). Factors involved in the synthesis of LPS and other lipids as well as the Tol-Pal system were also identified (Table S2), providing further support for the idea that *ibaG* is important for biogenesis and/or maintenance of the cell envelope.

**Figure 5.**
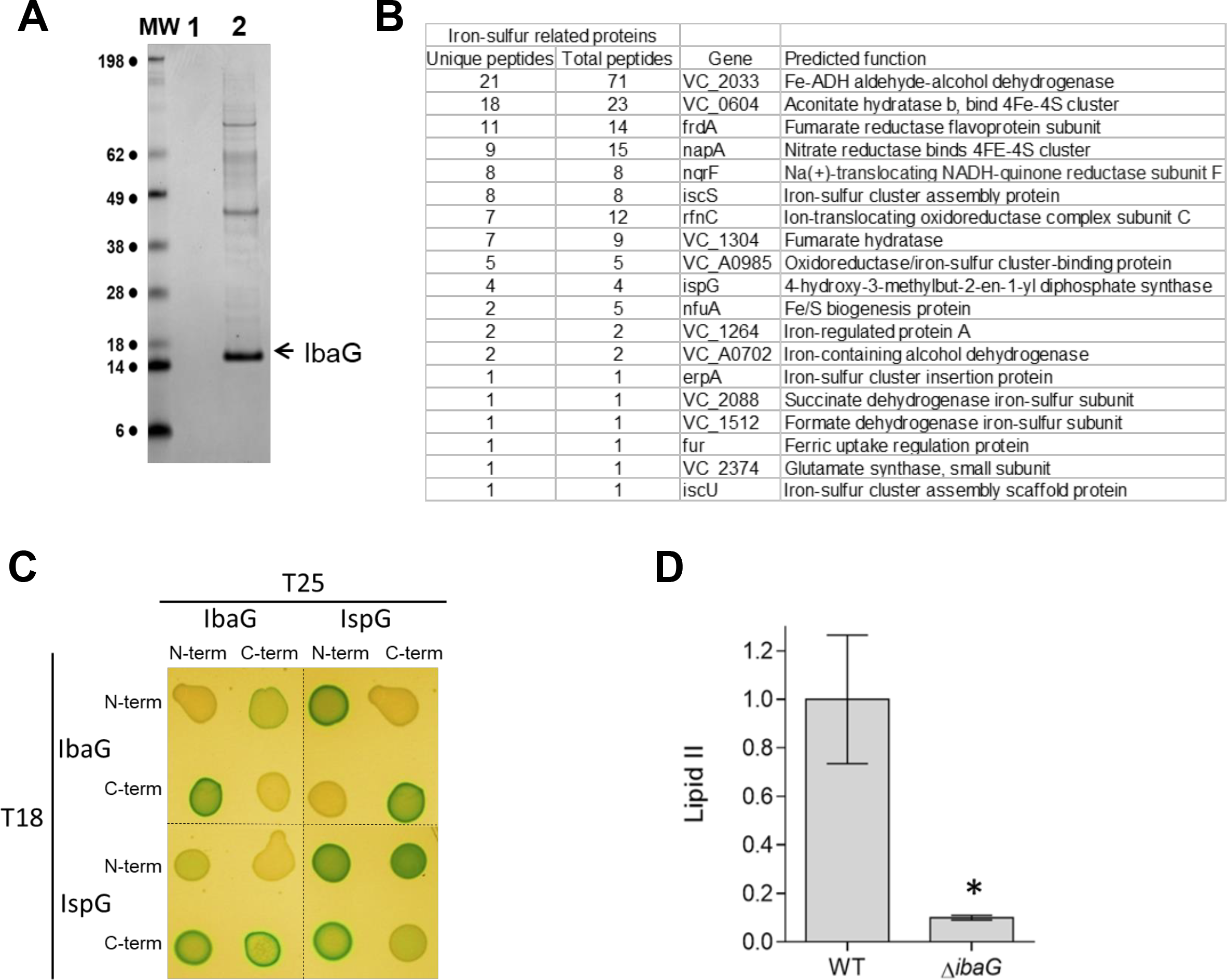
IbaG interacts with iron-sulfur cluster proteins. **A**. Coomassie Blue stained gel of proteins recovered after TAP purification from cell extracts of *V. cholerae* producing TAP-TAG only (1) or IbaG-TAP-TAG (2). Bands of interest were analyzed by mass spectrometry. **B**. IbaG-interacting proteins identified by mass spectrometry that are iron-sulfur containing proteins or facilitate biogenesis of iron-sulfur proteins (complete list of interacting proteins is presented in Table S2). **C**. Bacterial adenylate cyclase two-hybrid analysis of IbaG and IspG interactions. Colonies of *cya*-negative strains producing T25 and T18 fusions of the respective proteins on LB medium supplemented with X-Gal and IPTG are shown. **D**. Lipid II quantification in WT and Δ*ibaG* strain grown in M9 medium to exponential phase. * = p-value <0.01 (t-test).

IspG, one of the proteins that copurified with IbaG, contributes to the synthesis of precursors to lipid II, which mediates a critical early step in PG synthesis, suggesting a possible explanation for the reduced PG in the Δ*ibaG* cells. Bacterial two-hybrid analysis confirmed the interaction between IbaG and IspG (Fig. 5C). Furthermore, UPLC chromatography coupled to MS/MS analysis of Lipid II levels in exponential phase WT and Δ*ibaG* cells revealed markedly lower abundance (10-fold change) of Lipid II in the Δ*ibaG* cells (Fig. 5D). Reduced lipid II levels (and subsequent effects on PG synthesis and homeostasis) could also contribute to the Δ*ibaG* mutant’s increased sensitivity to antibiotics that target cell wall synthesis.

## Discussion

Here, we characterized the *V. cholerae* BolA-like protein IbaG. IbaG, which is encoded in the midst of loci that contribute to the biogenesis and maintenance of the cell envelope, likewise appears to modulate production and/or integrity of the *V. cholerae* envelope. Mutants lacking *ibaG* contain reduced amounts of peptidoglycan and LPS and have altered lipid profiles. Likely as a result of altered cellular barriers, ∆*ibaG V. cholerae* exhibit elevated sensitivity to antibiotics that target the cell wall and to detergents and other envelope-disrupting factors. The mutant also displays impaired capacity to colonize the intestine in an animal model of infection. Mutagenesis and biochemical analyses provided further support for the idea that *V. cholerae ibaG* contributes to cell envelope biogenesis and suggest that it may do so by modulating assembly and/or trafficking of iron-sulfur clusters.

Although *E. coli* and *V. cholerae ibaG* have significant homology and share genomic context, our findings revealed that deletion of *ibaG* has markedly distinct consequences in these two gamma proteobacteria. While no morphological defect was found for ∆*ibaG E. coli* (14), ∆*ibaG V. cholerae* were frequently elongated, branched, and wider than WT *V. cholerae.* Furthermore, *E. coli ibaG* is induced by acid stress and promotes survival in response to acid challenge (14), whereas neither phenotype was apparent in *V. cholerae*. Such diversity of function has previously been observed for a variety of *E. coli/ V. cholerae* homolog pairs involved in cell wall regulation (e.g., DacA-1/PBP5, PBP1A, AmiB, and NlpD) (24–26). Similarly, BolA has been found to play distinct roles in *Pseudomonas fluorescens* and *E. coli* (27), suggesting that each factor may be adapted to meet the specific needs of the host organism.

Interestingly, the *ibaG* mutant exhibited aberrant morphology during exponential phase growth, but normal size and shape during stationary phase. Growth in minimal vs rich media also exacerbated the mutant’s distorted morphology. It is possible that the increased demand for cell wall components associated with cell growth and division, coupled with the reduced levels of PG, the PG biosynthetic factor Lipid II, and LPS, may contribute to the *ibaG* mutant’s inability to maintain normal morphology during rapid growth. Potentially arguing against this hypothesis is the slower growth in minimal compared to LB media; furthermore, a recent analysis in *E. coli* revealed that nutrient limitation tended to reduce the effect of mutations on cell morphology (28). Given the apparent link between *ibaG* and Fe-S cluster–linked processes, it is possible that differences in iron availability in the minimal media contribute to the increased shape alterations rather than, or in addition to, the extent of nutrients.

In addition to its effect on cell morphology, the growth phase of the *ibaG* mutant also influenced its capacity to compete against WT *V. cholerae* in colonizing the intestine of a model animal host. When stationary phase cultures were used to infect infant rabbits, the *ibaG* mutant exhibited a less than two-fold deficit in colonization relative to the co-inoculated WT strain; in contrast, log phase cells exhibited an ~50-fold deficit. We speculate that replicating *ibaG* cells may be particularly sensitive to host protective factors that are encountered early in the infection process (e.g., bile), and therefore may be preferentially eliminated at the beginning of the infection process. Such a disadvantage is consistent with the mutant’s increased susceptibility to cell envelope-disrupting factors in vitro. The normal morphology of the *ibaG* cells in stationary phase may be indicative of a relatively unperturbed cell envelope that is more able to withstand such host defenses. Although the stationary phase inoculum gives rise to replicating (and presumably morphologically aberrant) cells in vivo, replication may occur after cells have reached intestinal sites where they are not exposed to high concentrations of agents such as bile.

Our analysis of proteins that co-purify with IbaG provided possible explanations for the reduced levels of cell envelope components observed in the ∆*ibaG* mutant. Several members of the RfB family, responsible for O-antigen synthesis, were found to interact directly or indirectly with IbaG; the absence of such interactions may contribute to ∆*ibaG V. cholerae’s* LPS deficiency. Similarly, an interaction between IbaG and IspG, which contributes to the synthesis of precursors to Lipid II, may underlie the reduction in Lipid II and PG that was observed in the *ibaG* mutant. Deficiencies in PG and LPS likely lead to formation of a cell wall and outer membrane that are defective in cell division, maintenance of turgor pressure and sensitive to membrane disrupting factors, accounting for some of the mutant’s phenotypes.

Notably, analysis of factors that co-purify with IbaG also suggest that *V. cholerae* IbaG is linked to production or trafficking of iron-sulfur clusters. We found that IbaG is able to interact directly or indirectly with several proteins involved in iron-sulfur biogenesis or containing iron-sulfur clusters, including IscU, IscS, and NfuA. Given the pivotal role of iron-sulfur containing proteins in numerous cellular processes, including central carbon metabolism, DNA/RNA metabolism, signal transduction, and stress responses, their interactions with IbaG suggest multiple ways that *ibaG* deletion might disrupt cellular physiology, which may account for its pleotropic effects. Finally, perhaps even more remarkable than the extreme distortion of the shape and size of exponential phase ∆*ibaG V. cholerae* is their capacity to regain normal shape and size; unraveling the mechanisms that enable this recovery should yield insight into the plasticity of bacterial shape determining pathways.

## Acknowledgments

We thank all the members of the Waldor lab for helpful discussions. Research in the MKW laboratory is supported by HHMI and NIH Grant R01 AI-042347. AZ was supported by an EMBO long-term fellowship (ALTF 1514-2016) and by a HHMI Fellowship of the Life Sciences Research Foundation. Research in the L.X laboratory is supported by NIH grant (R01 AI-136979). Research in the F.C lab was supported by the Knut and Alice Wallenberg Foundation (KAW), the Laboratory of Molecular Infection Medicine Sweden (MIMS), the Swedish Research Council and the Kempe Foundation.

## Material and Methods

### Strains, Media, and Growth conditions

All *V. cholerae* strains described in this study are derivatives of *V. cholerae* El Tor strain N16961 (29). *E. coli* DH5α λpir was used for general cloning purposes. *E. coli* SM10 λpir was used for conjugation. Cells were grown at 37°C in Luria-Bertani (LB) medium, or M9 medium supplemented with 0.2% glucose (M9). Media were supplemented when needed with 200 µg/ml streptomycin, 50 µg/ml carbenicillin (*V. cholerae*) or 20 µg/ml chloramphenicol (*E. coli)*. For induction of genes under control of arabinose-inducible promoters, strains were grown in media supplemented with 0.2% L-arabinose.

For growth curves, at least 3 replicates per strain and condition were grown in 200 µl medium in a 100 well honeycomb plate inoculated 1:100 from an exponentially growing pre-culture (OD_600_nm ~ 0.02) and analyzed in a BioScreen C growth plate reader at 10 min intervals. Data were analyzed using Microsoft Excel.

### Construction of plasmids and strains

Plasmids and strains are described in supplemental table S3. Plasmids were generated with Gibson assembly (30). In-frame deletions were introduced using sucrose-based counter selection with *sacB*-containing suicide vector pCVD442 (31). Proteins were overproduced by placing the respective gene under the control of the *araC* (P_BAD_) promoter using vector pBAD33 (32).

### Microscopy

Microscopy was performed using exponentially growing cells (OD_600nm_ ~ 0.2-0.4) or stationary phase cells. Bacteria were immobilized on 1% agarose pads, and were visualized using a Nikon Eclipse Ti equipped with a Andor NeoZyla camera and a 100× oil Phase3 1.4 NA objective. Images were processed using ImageJ (http://rsb.info.nih.gov/ij/) and MicrobeTracker (33) to generate cell length and width distribution histograms. The mean width, which is the average of the width over the entire cell, was measured instead of the maximum width, given the variation in width along *ibaG* cells. A non-parametric statistical analysis (Mann Whitney U test) was performed using Prism because of the non-normal distribution of cell sizes in the mutant strain (34). Staining with FM4-64 was performed as described (35). In brief, cells were grown to exponential phase or stationary phase in LB or M9 medium and 1 µg/mL of FM4-64 was added to the cultures and incubated for 5 min at room temperature and imaged as above.

### Bacterial two-hybrid assay

The adenylate cyclase-based bacterial two-hybrid technique was used as previously published (36). Briefly, IbaG, IspG and Grx4 were fused to the isolated T18 and T25 catalytic domains of the *Bordetella* adenylate cyclase. After transformation of the two plasmids producing the fusion proteins into the reporter BTH101 strain, plates were incubated at 30°C for 48 h. Three independent colonies for each transformation were inoculated into 600 μl of LB medium supplemented with ampicillin, kanamycin, and IPTG (0.5 mM). After overnight growth at 30°C. 10 μl of each culture were dropped onto LB plates supplemented with ampicillin, kanamycin, bromo-chloro-indolyl-galactopyrannoside (X-Gal 40 µg/mL) and IPTG (0.5 mM) and incubated for 16 h at 30°C. The experiments were done at least in triplicate, and a representative result is shown.

### Purification and visualization of lipopolysaccharide (LPS)

LPS was extracted following the protocol described in Davis and Goldberg (37). Briefly, pelleted bacteria harvested from exponential phase cultures were resuspended in 200 µl of SDS buffer (2% β-mercaptoethanol, 2% SDS and 10% glycerol in 0.05M Tris HCl, pH 6.8), and boiled for 15 min. The samples were treated with 5 µl of DNAse and RNAse (10 mg/mL) for 30 min at 37°C, then with 10 µL of Proteinase K (10 mg/mL) for 3 hours at 59°C. 200µL of ice-cold Tris-saturated phenol was then added and the samples were incubated 15 min at 65°C, with occasional vortexing. 1 mL of diethyl ether was added before centrifugation for 10 min at 20,600 x g and the bottom blue layer was extracted. The extractions with Tris-saturated phenol and diethyl ether were repeated twice before adding 2X SDS-buffer to the samples. 15 µl of samples were run on SDS-polyacrylamide gels. LPS was visualized using the Pro-Q Emerald 300 Lipopolysaccharide Gel Stain Kit (Molecular Probes) according to the manufacturer’s instructions.

### Acid resistance assay

Bacterial cultures were grown in LB until OD_600nm_~ 0.3, then diluted 20-fold in LB pH5.5 and incubated for one-hour before plating of serial dilutions to determine CFU/mL for each strain. CFU/mL were similarly determined for growth in LB pH7 prior the acid challenge and the relative survival (CFU/mL pH5.5 / CFU/mL pH7) was calculated to determine acid resistance for both strains. The pH of the LB broth was adjusted using 1 mM HCl.

### Quantitative-PCR

Cells from overnight (stationary phase) cultures were inoculated in triplicate into 5 ml LB or M9, grown at 37°C until exponential phase (OD_600nm_ ~ 0.3) or stationary phase. Total RNA was extracted from harvested cells with TRIzol reagent (Life Technologies). RNA was treated with Turbo DNase I for 30 min (Life Technologies) and subjected to qRT-PCR as previously described (38). Briefly, 1 μg total RNA was used for the reverse transcription reaction with Superscript III first strand synthesis system with random hexamers (Life Technologies). Real time-PCR amplification of the synthesized cDNA was conducted using the Fast SYBR Green Master Mix kit (Life Technologies). Each of the three biological replicates was analyzed in technical triplicate on the StepOnePlus platform (Life Technologies) using primers shown in supplemental table 3. The data was analyzed by ΔΔCT method using *rpoC* mRNA as an internal control. Log 2 (Fold change) was calculated from ΔΔCT results.

### Transposon mutant library construction and sequencing

Transposon insertion sequencing was performed as described previously (39). Transposon libraries were created in WT and Δ*ibaG V*. *cholerae* using the transposon delivery vector pSC189. ~600,000 transposon mutants were generated for each strain. Genomic DNA was purified and sequenced on an Illumina MiSeq benchtop sequencer (Illumina, San Diego, CA). Sequenced reads were mapped onto the N16961 *V.cholerae* reference genome, and all TA sites were tallied and assigned to annotated genes as previously described (40). Insertion sites were identified as described (39), and significance was determined using the Con-Artist pipeline.

### MIC assay

MIC assays were performed using an adaptation of a standard methodology with exponential-phase cultures (41). In short, serial 2-fold dilutions of the antimicrobial agents were prepared in 50 µl of LB in a 96-well plates. Then, to each well was added 50 µl of a culture prepared by diluting an overnight culture 1,000-fold into fresh LB broth, growing it for 1 h at 37°C, and again diluting it 1,000-fold into fresh medium. The plates were then incubated without shaking for 24 h at 37°C.

### Peptidoglycan (PG) purification and analysis

PG samples were prepared and analyzed in triplicates as described previously (42, 43). Briefly, 1 L of exponential WT and ∆*ibaG* grown in LB were harvested and boiled in 5% SDS for 2 h. Sacculi were repeatedly washed with MilliQ water by ultracentrifugation (110,000 rpm, 10 min, 20ºC) until total removal of the detergent, followed by digestion with pronase E (100 ug/ml) for 1h at 60ºC. Finally, samples were treated with muramidase (100 μg/ml) for 16 hours at 37ºC. Muramidase digestion was stopped by boiling and coagulated proteins were removed by centrifugation (10 min, 14,000 rpm). For sample reduction, the pH of the supernatants was adjusted to pH 8.5-9.0 with sodium borate buffer and sodium borohydride was added to a final concentration of 10 mg/mL. After incubating for 30 min at room temperature, the samples pH was adjusted to pH 3.5 with orthophosphoric acid.

UPLC analyses of muropeptides were performed on a Waters UPLC system (Waters Corporation, USA) equipped with an ACQUITY UPLC BEH C18 Column, 130Å, 1.7 μm, 2.1 mm X 150 mm (Waters, USA) and a dual wavelength absorbance detector. Elution of muropeptides was detected at 204 nm. Muropeptides were separated at 35ºC using a linear gradient from buffer A (phosphate buffer 50 mM pH 4.35) to buffer B (phosphate buffer 50 mM pH 4.95 methanol 15% (v/v)) in a 20 minutes run, with a 0.25 ml/min flow.

Relative total PG amounts were calculated by comparison of the total intensities of the chromatograms (total area) from three biological replicas normalized to the same OD600 and extracted with the same volumes. Muropeptide identity was confirmed by MS/MS analysis, using a Xevo G2-XS QTof system (Waters Corporation, USA). Quantification of muropeptides was based on their relative abundances (relative area of the corresponding peak) normalized to their molar ratio.

### *In vivo* colonization assay

Intestinal colonization in infant mice was carried out as described previously (44). Cells for the exponential phase inoculum were grown separately to OD_600nm_ ∼ 0.3 in LB, then diluted 1:100 in the same medium prior to mixing 1:1. Cells for the stationary phase inoculum were grown separately overnight in LB at 37°C, then diluted 1:1000 in LB prior to mixing. Infant mice were gavaged with 50 μL of the 1:1 inoculum mixture, then sacrificed after 24 hours. Dilutions of homogenized small intestines were plated on LB agar to enumerate CFU. Competitive indices (CI) were calculated as the ratio of mutant to WT bacteria isolated from intestines normalized to the input ratio. Statistical significance was determined using a two-tailed Mann-Whitney U *t*-test (p-value < 0.01).

### Lipid II quantification

Precursor extraction was performed as described previously and performed in triplicates (45). Briefly, 500 ml of WT and ∆*ibaG* were grown in LB to OD600 0.45. Cells were harvested, resuspended in 5 ml PBS in 50 ml flasks containing 20 ml CHCl3: Methanol (1:2). The mixture was stirred for 1h at room temperature and centrifuged for 10 min at 4000 x g at 4ºC. The supernatant was transferred to 250 ml flasks containing 12 ml CHCl3 and 9 ml PBS, stirred for 1h at room temperature and centrifuged for 10 min at 4000 x g at 4ºC. The interface fraction (between the top aqueous and bottom organic layers) was collected and vacuum dried. To remove the lipid tail, samples were resuspended in 100 μl DMSO, 800 μl H2O, and 100 μl ammonium acetate (100 mM pH 4.2). This mixture was boiled for 30 min, dried in a vacuum and resuspended in 300 μl H2O.

Samples were analyzed by UPLC chromatography coupled to MS/MS analysis, using a Xevo G2-XS QTof system (Waters Corporation, USA). Precursors were separated at 45°C using a linear gradient from buffer A (formic acid 0.1% in water) to buffer B (formic acid 0.1% in acetonitrile) in an 18-minute run, with a 0.25 ml/min flow. A library of compounds was used to target the identification of peptidoglycan precursors and possible intermediates, although only lipid II was detected. Lipid II amounts were calculated based on the integration of the peaks (total area), normalized to the culture OD.

### Tandem affinity purification assay and Mass Spectrometry analysis

IbaG was purified using a standard TAP protocol. Briefly, an overnight culture of *V. cholerae* encoding IbaG C-terminally tagged with calmodulin binding protein, a TEV cleavage site and protein A was used to inoculate 500ml of LB (1/100 dilution), which was grown 5 h 30 at 37°C with shaking and then cells were pelleted and washed in cold PBS. Tandem affinity purification was then performed as described before (46, 47). Then, the cells were broken using an Emulsilfex C3 (Avestin) in presence of proteases inhibitor (Complete, Roche). The lysate was used to bind to 200ul of IgG Sepharose beads (Amersham Biosciences) for 2h at 4°C using a disposable chromatography column (BioRad). The IgG-Sepharose column was washed with 35 ml of protein A binding buffer (10 mM Tris–HCl, pH 8, 150 mM NaCl, 0.1% NP-40), followed by 10 ml of the TEV cleavage buffer (10 mM Tris–HCl, pH 8, 150 mM NaCl, 0.1% NP-40, 0.5 mM EDTA, 1 mM dithiothreitol DTT). Cleavage with TEV was performed using 10 ul (100 units) of AcTEV (Invitrogen) in 1ml of cleavage buffer for 2h at 4°C. Calmodulin-Sepharose (Stratagene) purification was performed as described (47). Independent tandem affinity purifications followed by mass spectrometry analysis was performed at least twice.

### Homology alignments and structural predictions

The 3D homology model of BolA and IbaG from *V. cholerae* were constructed using the Phyre2 Server (48) (www.sbg.bio.ic.ac.uk/~phyre2). The program used BolA (PDB id: 2DHM) and YrbA (PDB id: 1NY8) from E. coli as template to generate the models. PyMOL (The PyMOL Molecular Graphics System, Version 1.2r3pre, Schrödinger, LLC.) was used to generate the figure. Multiple sequence alignments were assembled from selected pairwise alignments and converted to clustal format (49) and uploaded to Ali2D (http://toolkit.tuebingen.mpg.de/ali2d) to generate images for secondary structure similarity (50).

### Lipid quantification

Extraction of lipids from *V. cholerae* pellets was performed using the method of Bligh & Dyer, as described previously (51–54). Briefly, dried pellets (Δ*ibaG*: 74.1 ± 6.0 mg; N16961: 71.9 ± 4.2 mg; t-test *P* = 0.62) in 10 mL glass centrifuge tubes (Fisher) were reconstituted in 1 mL of H_2_O (Fisher Optima LC-MS) and sonicated for 30 min in an ice bath, followed by the addition of 4 mL of chilled 1:2 chloroform/methanol (Fisher Optima LC-MS) extraction solution. Following mixing and centrifugation, the organic phase of the two-layer extraction was collected into fresh glass centrifuge tubes and dried in a vacuum concentrator. Extracts were reconstituted with 500 µL of 1:1 chloroform/methanol solution. For analysis, 5 µL of extract was transferred to an LC vial, dried, and reconstituted with 100 µL of 2:1 acetonitrile/methanol solution. A pooled quality control sample was prepared from 15 µL of each sample.

Characterization of the *V. cholerae* lipidome was performed by hydrophilic interaction liquid chromatography (HILIC) coupled to ion mobility-mass spectrometry (IM-MS), as described previously (51). Data was acquired for each sample in both positive and negative electrospray ionization modes over the range of 50-1200 *m/z*. Alignment of HILIC-IM-MS data and peak detection were performed in Progenesis QI (Nonlinear Dynamics) with the default “All Compounds” normalization method. The negative mode dataset was filtered by ANOVA P ≤ 0.1, which retained 528 features. The top 10 features for phosphatidylethanolamines (PEs) and phosphatidylglycerols (PGlys) and the top 5 features for lyso-phosphatidylethanolamines (L-PEs) and lyso-phosphatidylglycerols (L-PGlys) that meet the ANOVA P threshold were summed in the figure. Student’s t-tests for two samples were performed using a two-tailed distribution and equal variance. Identification of lipid species was performed against the METLIN database within 15 ppm mass accuracy (55, 56).

---

**Figure S1. Comparison of secondary structures and amino-acid sequences of BolA and IbaG in *E. coli* and *V. cholerae***

**A-C.** ClustalW alignment of predicted amino-acid sequences of BolA and IbaG from *E. coli* (E.c) and *V. cholerae* (V.c). The KEGG database was used to obtain the protein sequences which were aligned using ClustalW NPSA.

**D.** Clustal Omega alignment of BolA and IbaG from *V. cholerae*. Prediction of secondary structures were generated using Ali2D and PSIpred. Alpha helices (H) and beta-sheets (E) are colored in red and blue, respectively. The helix turn helix motif (HTH) is annotated.

**Figure S2. Morphology and growth of *bolA* and *ibaG* mutant cells**

**A**. Phase contrast and fluorescence imaging of FM4-64 stained Δ*bolA* and *bolA* overexpressing (bolA++) cells grown to exponential phase and stationary phase in M9 medium. Scale bars 2 μm.

**B**. Phase contrast and fluorescence imaging of FM4-64 stained Δ*ibaG* cells grown to exponential phase in M9 and LB media. Scale bars 2 μm.

**C**. Cell length and mean width distribution of WT and Δ*ibaG* strains grown to stationary phase in M9 medium. At least 1000 cells were measured for each condition using MicrobeTracker; the differences in distributions of lengths and widths (determined with a Mann Whitney test) were not significant (p-value >0.15 for both).

**Figure S3. Complementation and *ibaG* expression analysis**

**A**. *IbaG* expression in WT V. cholerae in different conditions measured by quantitative-PCR. WT cells were grown in LB until exponential phase (OD_600nm_~ 0.3), then diluted 20-fold in LB pH5.5 or LB pH7 and grown for one hour before RNA samples were processed for qPCR. WT cells were also grown in LB and M9 until exponential phase or stationary phase before processing samples for qPCR. The data was analyzed by ΔΔCT method using *rpoC* mRNA as internal control. Log 2 (Fold change) was calculated from ΔΔCT results. The reference is expression of *ibaG* in LB pH7. Experiments were performed with biological triplicates and the standard deviation is shown.

**B**. Phase contrast and fluorescence imaging of FM4-64 stained Δ*ibaG* cells complemented with *ibaG* expressed from plasmid pBAD33. Cells were grown to exponential phase in M9 medium supplemented with 0.2% arabinose. Scale bars 2 μm.

**C**. Growth curves of indicated strains cells grown in M9 medium supplemented with 0.2% arabinose. OD_600nm_ was measured at 10-min intervals. Experiments were done in biological triplicate; standard deviations are shown.

**D**. Growth curves of WT and Δ*ibaG* strains grown in LB pH7 and LB pH5.5. OD_600nm_ was measured at 10-min intervals. Experiments were performed in triplicate; error bars show standard deviations.

**Figure S4. Chromatograms of peptidoglycan analysis**

Chromatograms from analysis of peptidoglycan derived from WT and ∆*ibaG* strains in exponential phase (panel A). The muropeptides identified in each peak are described in the table (panel B).

**Figure S5. Interaction between IbaG and Grx4**

Bacterial adenylate cyclase two-hybrid analysis of IbaG and Grx4 interactions. Colonies of *cya*-negative strains producing T25 and T18 fusions of the respective proteins on LB medium supplemented with X-Gal and IPTG are shown.

**Supplemental Tables**

**Table S1. Transposon-insertion sequencing analysis**

Comprehensive list of the genes over or under-represented in the Δ*ibaG* library with a mean fold change >2 and a p-value <0.05.

**Table S2. Total proteins identified by mass spectrometry**

List of the proteins identified by mass spectrometry and their functional classification represented on a pie chart (numbers represent the percentage of genes in each category).

**Table S3. Strains, plasmids and oligos**

Strains, plasmids and oligos used in this study.

